# Atlas of Cynomolgus Macaque Hematopoiesis

**DOI:** 10.1101/2024.04.22.590220

**Authors:** Shinichiro Oshima, Rahul Sinha, Kazuhiro Takeuchi, Ken Mizuta, Tomonori Nakamura, Ryoji Ito, Ikuo Kawamoto, Chizuru Iwatani, Masataka Nakaya, Hideaki Tsuchiya, Shruti Bhagat, Tomoyuki Tsukiyama, Akifumi Takaori-Kondo, Mitinori Saitou, Yasuhiro Murakawa, Ryo Yamamoto

## Abstract

Self-renewal and differentiation are inherent properties of hematopoietic stem cells (HSCs) that are necessary to support hematopoiesis; however, the underlying mechanisms, especially in human, remain unclear. Here, using the cynomolgus macaque as a surrogate model, we develop a new gating strategy to isolate with high purity transplantable cynomolgus HSCs and generated a single-cell transcriptomic map of cynomolgus HSCs and progenitor cells — that covering gestational periods previously not analyzed in human. We show that hematopoietic cells from the late-1^st^ to early-3^rd^ trimester fetal liver and late-2^nd^ trimester and thereafter bone marrow have repopulating potential, closely mimicking humans. Unexpectedly however, we found unlike in human, cynomolgus HSCs express CD38 but not CD33, indicating that these cellular counterparts are molecularly distinct. Our transcriptomic analysis reveals the presence of a direct differentiation pathway from HSCs to megakaryocyte lineages, lineage-primed multipotent progenitors and also identified putative HSC surface markers. Taken together, our comprehensive dataset highlights not only the utility of cynomolgus monkeys as model systems to study hematopoiesis but also their potential for translational applications.

## Introduction

Hematopoietic stem cells (HSCs) are characterized by their ability to self-renew and differentiate into multiple lineages, playing a vital role in sustaining hematopoiesis to ensure the production of mature blood cells throughout the lifespan of an individual. To experimentally validate such properties/features of HSCs, to-date, the gold-standard approach has been to successful reconstitute hematopoiesis in the recipient using purified donor HSCs, thereby requiring animal model systems for transplantation/xenograft assays^1^. To this extent, the mouse has served as an excellent model system to better understand HSCs during development^2^ and to identify candidate markers that can be used to purify HSCs. Despite similarities between the mouse and human hematopoietic systems, such as the principle hematopoietic sites including the fetal liver (FL), fetal bone marrow (FBM), and various other anatomical sites^2^ during embryogenesis — fundamental differences exist— including immune tolerance, anatomy of extraembryonic tissues, pregnancy duration and timing of birth, and species-specific regulatory mechanisms^3,4,5,6,7^, which prevent the direct transfer and translational application of knowledge between mouse and human, especially in the clinical setting. Though several studies have reported the functionality and molecular features of human fetal hematopoiesis^8,9,10,11,12,13^, due to the limited accessibility and ethical constraints associated with the use of human fetal tissues^14^, such research has been restricted to early phases of human fetal development. Therefore, in order to establish a better understanding of hematopoiesis and HSC development in humans, orthologous systems that more closely recapitulate events occurring in human are necessary.

Being the closest relatives to human —nonhuman primates (NHPs) are a powerful surrogate model system to investigate processes that either completely or partially recapitulate those in human, including fetal development and pathogenesis of diseases. To date, NHPs are commonly used in preclinical studies such as for pharmaceutical development and in gene therapy^15,16,17^. One of the selective advantages of using NHPs lies in the ability to obtain fetal samples on precisely targeted days across wide range of pregnancy stages. Furthermore, NHPs are also amenable to autologous bone marrow transplantation, as previously shown in rhesus monkey *(Macaca mulatta)* and baboon^18^, indicating their potential in studying HSCs and fetal hematopoiesis. Despite the human HSC marker CD34 having been previously used as a common stem cell marker in NHPs ^19,20,21,22^, whether even CD34 is a bona fide marker for all NHP HSCs remains unclear. Thus, a better characterization of hematopoietic stem and progenitor cells (HSPCs) in NHPs is necessary in order to establish definitive HSC markers and to distinguish the similarities and differences in hematopoietic systems between humans and NHPs. This will then also further clarify how NHP systems can be properly used in preclinical settings.

Among NHPs, rhesus monkeys are commonly used as model systems, especially in the field of viral pathology — though no evidence demonstrating any selective advantages over other NHPs have been described ^23,24,25,26,27^. In fact, with current pressures on rhesus stocks as a result of expanding COVID-19 research^28,29^ and global social and political situations, as well as the limitations associated with defined breeding seasons and inability to give birth all year around, this suggests the potential of use of other monkeys as model systems to investigate developmental processes including hematopoiesis. In this respect, cynomolgus monkeys *(Macaca fascicularis)* are thought to be a next model for such research purposes due to their ability to give birth all year round without distinct breeding season — similar to humans — allowing easier access to fetal samples.

Here, using the cynomolgus monkey as a model system, we investigate the heterogeneity of HSPCs across multiple developmental stages and tissues. We establish a new gating strategy to purify transplantable cynomolgus HSCs and construct a single-cell transcriptomic map of cynomolgus hematopoiesis, which we anticipate will become useful resource for future preclinical studies using cynomolgus monkey HSPCs.

## Results

### Cynomolgus fetal liver and fetal bone marrow has repopulating potential in peripheral blood

The average gestation length of cynomolgus macaque is 165 days and consists of the 1^st^ (day 0-55), 2^nd^ (day 56-110) and 3^rd^ trimesters (day 111-165). We isolated samples during embryonic development from presumed hematopoietic sites at embryonic day (E) 49 (late 1^st^ trimester); E57-E61 (early 2^nd^ trimester); E96-E96 (late 2^nd^ trimester); E121 (early 3^rd^ trimester); and adult (9-13-year-old) (**Figure 1A**). To determine whether transplantable HSCs are present in cynomolgus FL and FBM, we performed xenotransplantation assay in mice. Un-fractionated whole cells (E49-FL and E58-FBM) or CD34^+^ cells purified by magnetic cell separation (MACS) (E58-FL, E96-FL, E96-FBM, E121-FL, E121-FBM) were transplanted into NSG mice after sub-lethal dose irradiation (**Figure 1B**). Analyzing the peripheral blood (PB) of recipient mice every 4 weeks after transplantation up to 16 weeks post-transplant, we found mature donor cynomolgus CD45^+^ blood cells were not detected in recipient mice transplanted with E58-FBM cells. In contrast, we observed successful reconstitution in PB using FL (E49, E58, E96, E121) or FBM (E96, E121). These results indicate that transplantable HSCs are present within the FL (from E49-E121) and FBM (from E96-E121) that have capacity to repopulate blood cells in immunodeficient mice and that cynomolgus HSC transition from FL to FBM occurs during the 2^nd^ trimester, consistent with human^12^. Furthermore, though the exact disappearance of transplantable HSCs in human FL remains unclear, our results clearly show that transplantable HSCs are still present in the cynomolgus FL at early 3^rd^ trimester.

**Figure 1;.**
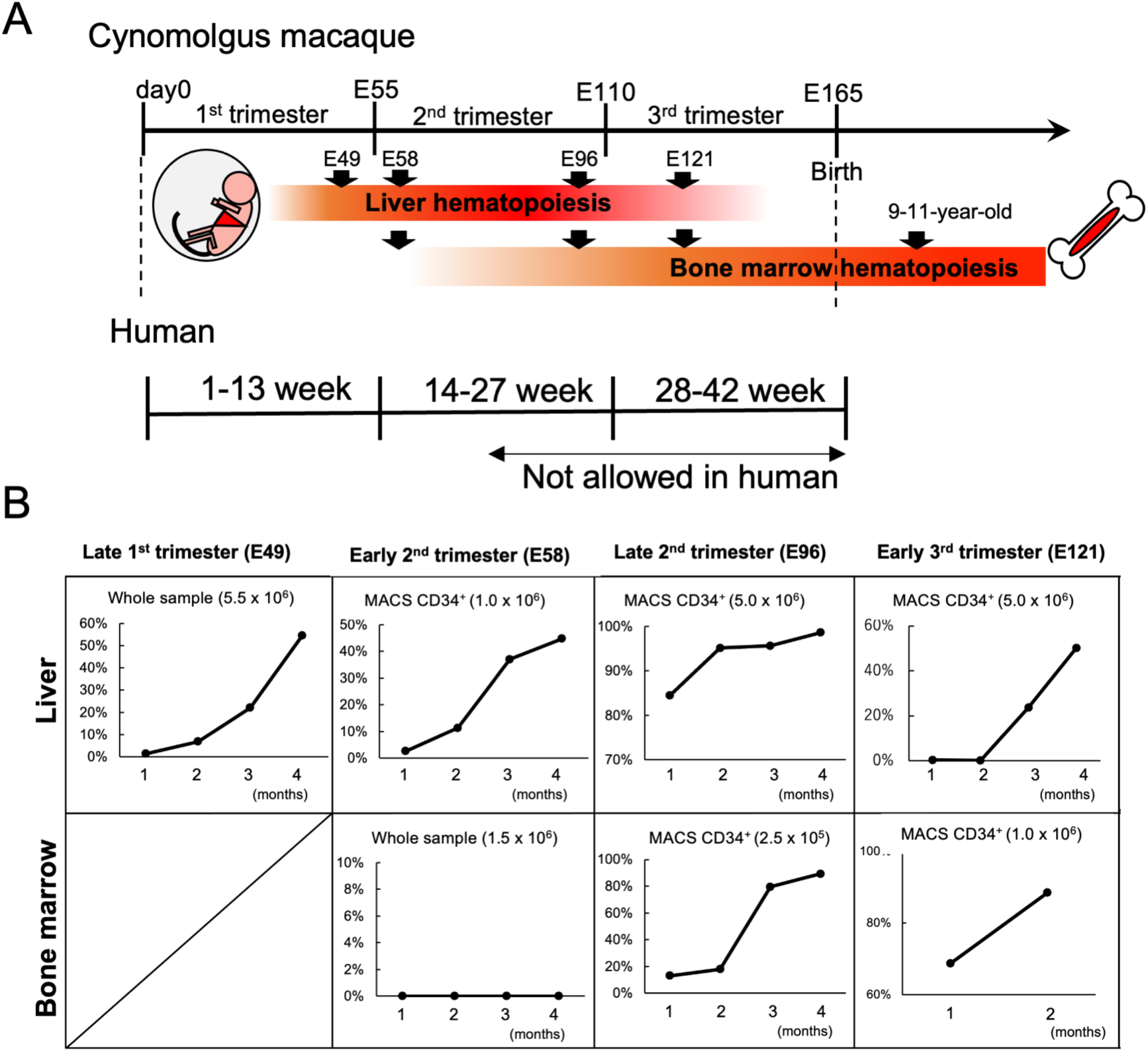
Identification of transplantable HSCs across multiple developmental stages and tissues. A: Schematic overview of the fetus and adult sampling. FL cells of late-1^st^ to early-3^rd^ trimester, and FBM cells of early-2^nd^ trimester to early-3^rd^ trimester were sampled. The red gradient bar shows presumed hematopoiesis across developmental stages. B: % Chimerism of cynomolgus CD45^+^ cells in recipient peripheral blood at indicated time point (months) after transplantation. Whole late 1^st^ FL or early 2^nd^ FBM cells or MACS-purified CD34^+^ early 2^nd^ FL, late 2^nd^ FL and FBM, or early 3^rd^ FL and FBM with indicated cell number were transplanted into NSG mice with 2.5-2.8 Gy irradiation.

### Purification and functional characterization of cynomolgus HSPCs

In cynomolgus monkeys, with the exception of CD34, no surface markers have been experimentally validated using the transplantation setting^30^. To determine the optimal combination of surface markers for isolating highly-purified transplantable HSCs, we first utilized surface markers that were established to purify HSCs in rhesus and pigtail macaques: Lineage (Lin)^-^CD45^mid^CD34^hi^CD45RA^-^CD90^+ 31^. Using these criteria, we isolated immunophenotypically-defined, distinct populations of presumable HSCs enriched CD45RA^-^CD90^+^ cells (Fraction 2: Fr2) and remaining the CD45RA^+^ or CD90^-^ cells (Fraction 3: Fr3) within Lin^-^CD45^mid^CD34^hi^ cells (Fraction 1: Fr1) for each developmental stage/tissue (**Figure 2A**). We noted that the frequency of Fr1 decreases with age in both FL and FBM (**Figure S1A**) and the frequency of Fr2 in late 2^nd^ trimester is highest among FL (from early 2^nd^ to late 3^rd^) or FBM (from late 2^nd^ to adult) (**Figure 2B**).

To purify transplantable HSCs among Fr2, we further selected several additional flow cytometric candidate antibodies and subsequently performed transplantation assays. The selection criteria used for antibodies were as follows: (1) those that are commonly used for human HSPC isolation, (2) those which are commercially available, and (3) those that have been previously reported to react with cynomolgus cells^32,33^. We tested CD11b (Mac-1), which is a marker for myeloid lineages; CD123, which is a marker for granulocytic monocytic progenitors (GMPs); CD127, which is a marker for common lymphoid progenitors (CLPs); CD71, which is a marker for erythroid lineages; CD41a, which is a marker for megakaryocyte lineages; CD31, which is a marker for endothelial cells; and CD33, and CD38, which are markers for HSPCs. There were few CD11b-, CD123-, CD127- or CD71-positive cells in Fr2 for each developmental stage/tissue, and partially positive for CD41a (**Figure 2C**). Surprisingly, we found that the majority of Fr2 is negative for CD33 and positive for CD38, which is the opposite to known human HSC expression patterns^34,35^. To clarify these discrepancies, we first fractionated Lin^-^CD45^+^ cells of early 2^nd^ FL into 4 fractions: CD34^low^CD38^-^, CD34^low^CD38^+^, CD34^hi^CD38^-^, and CD34^hi^CD38^+^. We found that only the CD34^hi^CD38^+^ cells had repopulating capabilities in NOG-W41 mice, while CD34^hi^CD38^-^ cells —which is typically used to purify human HSCs— had no repopulating potential (**Table 1, Figures S1B, S1C and S1D**). Next, in order to verify whether rhesus/pigtail macaque scheme can be applied to cynomolgus monkeys, we transplanted Fr2 and Fr3 cells of early 2^nd^ FL. Only Fr2 were found to have repopulating activity in NOG-W41 (**Table 1 and Figure S1D**). Furthermore, this gating scheme was also applicable to cynomolgus adult bone marrow (ABM) (**Figure S1D**). Finally, we found that CD33^+^ cells among Fr2 could not be engrafted (**Table 1**), indicating indeed, that cynomolgus HSCs are negative for CD33, which is the opposite to human. Furthermore, as CD33 expression progressively decreases with development in Fr2, this suggests that CD33 negativity can also be used to purify HSCs especially in early developmental stage (**Figure 2C**).

**Figure 2;.**
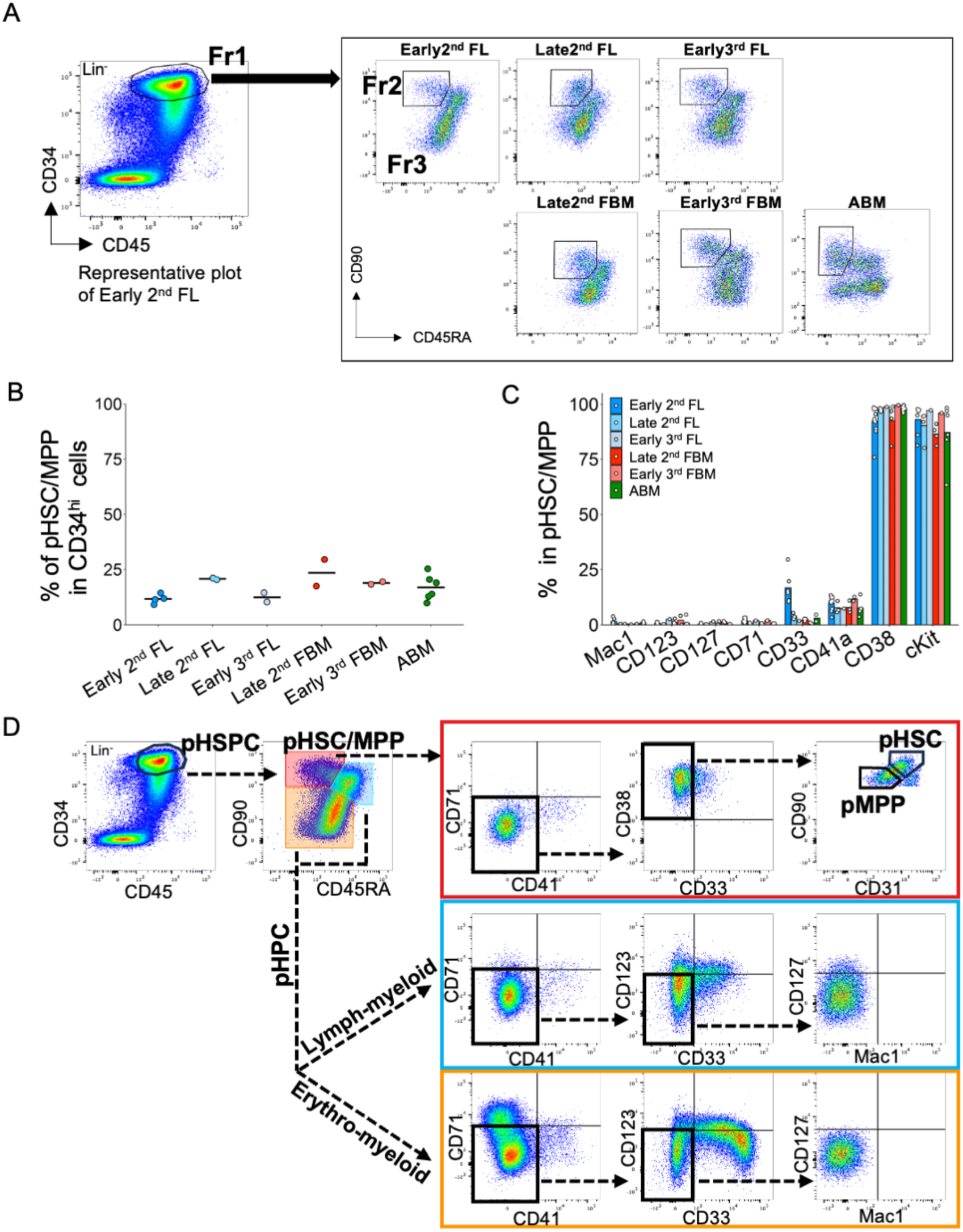
Functional purification of HSCs. A: Rhesus/pigtail macaque gating scheme reported by Radtke et al. was adopted to each developmental stage/tissue; CD45^mid^CD34^hi^ (Fraction 1, Fr1), CD45^mid^CD34^hi^CD45RA^-^CD90^+^ (Fraction 2, Fr2), and CD45^mid^CD34^hi^CD90^-^ or CD45^mid^CD34^hi^CD45RA^+^CD90^+^ (Fraction 3, Fr3). The number of cells for CD34^hi^ were normalized to 10,000 cells. B: The frequency of Fr2 for each developmental stage/tissue. The bars indicate the average value of the frequency. C: The frequency of Mac1, CD123, CD127, CD71, CD33, CD41a, or CD38-positive cells among Fr2. Each dot represents each sample. Each bar means each average of the frequency. D: A updated flow cytometric gating strategy to purify HSCs and to fractionate HSPCs. Representative flow cytometry dot data were shown using early 2^nd^ FL cells. pHSC/MPP (Lin^-^ CD45^+^CD34^hi^ CD45RA^-^CD90^+^), pHSC (Lin^-^CD45^+^CD34^hi^CD41^-^CD71^-^CD33^-^CD38^+^CD45RA^-^ CD90^hi^CD31^hi^), pMPP (Lin^-^CD45^+^CD34^hi^CD41^-^CD71^-^CD33^-^CD38^+^CD45RA^-^CD90^mid^CD31^mid^), lymph-myeloid pHPC (Lin^-^CD45^+^CD34^hi^CD45RA^+^CD90^+^), and erythro-myeloid pHPC (Lin^-^ CD45^+^CD34^h^ CD45RA^-^CD90^-^). The gate colors on CD45RA vs CD90 plot corresponds to those of box-outlines gating on downstream plots.

**Table 1;.** Functional purification of early 2nd FL-HSCs. Summary of transplantation assay for purifying transplantable HSCs. Freshly isolated and sort-purified cells, derived from early 2^nd^ FL, were injected in indicated quantities into NOG-W41 mice. Indicated transplanted populations were among live, lineage^-^ and CD45^+^ population. Engrafted mice (cynomolgus CD45^+^ cell greater than 0.1% at 3 months post-transplantation) out of total transplanted mice are listed.

Next, to refine further the purity of transplantable HSCs, we fractioned for CD31^hi^CD90^hi^ in Fr2 cells. CD31 is a well-known vascular endothelial cell marker ^36,37^ and is also present at a lower level on the surface of mouse hematopoietic cells including HSCs^38^. Based on expression pattern of CD31 and CD90 (**Figure 2D**), we compared the repopulating potential of CD31^mid^CD90^mid^ vs CD31^hi^CD90^hi^ in CD34^hi^CD45RA^-^CD90^+^CD41a^-^CD33^-^ population **(Table1)**. We found that CD31^hi^CD90^hi^ have repopulating activity, while CD31^mid^CD90^mid^ fraction did not and therefore we named these two populations pHSC and pMPP, respectively. Additionally, we also redefined Fr1 as pHSPC, Fr2 as pHSC/MPP, and Fr3 as pHPC (**Figure 2D**).

Having established our new gating strategy, we purified transplantable HSCs (estimated at 1 in 6204 cells) in early 2^nd^ FL-pHSC, which is 2.65-fold higher in purity compared to originally implemented gating schemes for rhesus/pigtail macaques (1 in 16,494 cells in pHSC/MPP) (**Figure S1E and Table S1**). Interestingly, the estimated frequency of transplantable HSCs was even higher in late 2^nd^ FL, at 1 in 1888 cells in pHSC/MPP, suggesting that HSCs expand in FL across early to late 2^nd^ trimester (**Figure S1E and Table S1**). Furthermore, we verified that both early 2^nd^ FL and ABM contain cells capable of reconstituting secondary recipients, suggesting that pHSC/MPP have long-term self-renewal capacity **(Figure S2)**.

We then analyzed surface expression patterns of progenitor cells and performed *in vitro* colony assay for further functional characterization. (**Figures 2D, S3A and S3B**). In pHPC, we detected two populations; relatively CD45RA high and CD90 high population (defined as CD45RA^+^CD90^+^ pHPC) and relatively CD45RA low and CD90 low population (defined as CD45RA^-^CD90^-^ pHPC). We found that CD45RA^+^CD90^+^ pHPC was the only fraction expressing CLP marker (CD127), while also being positive for GMP markers (CD123, CD33), but with no erythroid marker (CD71) expression. On the other hand, CD45RA^-^CD90^-^ pHPC expressed erythroid marker and GMP markers. Consistently, colony formation assay revealed that both these two populations could generate myeloid colonies, and that CD45RA^-^CD90^-^ pHPC gave rise to more erythroid colonies than CD45RA^+^CD90^+^ pHPC, indicating that CD45RA^+^CD90^+^ pHPC are lymphoid-myeloid pHPC and CD45RA^-^CD90^-^ pHPC are erythroid-myeloid pHPC. In all pHSC/MPP subfractions, both erythroid and myeloid colonies were detected. However, GEMM colonies were only detected in CD41^-^CD71^-^CD33^+^ pHSC/MPP or pHSC (CD41^-^CD71^-^CD33^-^ CD31^hi^CD90^hi^pHSC/MPP), suggesting CD90^hi^ could be the source of multipotent cells.

Taken together, these findings demonstrate that the most primitive HSCs were enriched in Lin^-^ CD45^mid^CD34^hi^CD45RA^-^CD41^-^CD33^-^CD90^hi^CD31^hi^ cells (pHSC), whereas lympho-myeloid progenitor cells were enriched in CD45RA^+^CD90^+^ subset of pHPC and erythro-myeloid progenitor cell were enriched in CD45RA^-^CD90^-^ subset of pHPC **(Figures 2D and S3C)**.

### Single-cell transcriptomic landscape of cynomolgus CD34^hi^ hematopoietic cells of FL, FBM and ABM

To determine the transcriptomic heterogeneity of cynomolgus hematopoiesis, we performed 10X Genomics 5’ chemistry based single-cell RNA-sequencing (scRNA-seq) data analysis of broad spectrum of hematopoietic cells, which were isolated using fluorescence-activated cell sorting (FACS) for Lin^-^ CD45^+^CD34^low^ (CD34^low^), pHSC/MPP and pHPC from early 2^nd^ FL **(Figure S4A)**. The Uniform Manifold Approximation and Projection (UMAP) integration revealed that CD34^low^ cells were primarily composed of mature cells, while pHSC/MPP and pHPC were composed of various types of immature HSPCs. (**Figures S4B and S4C**).

To further define the molecular characteristics of HSPCs across developmental stages, we analyzed CD34^hi^ hematopoietic cells of early 2^nd^ FL, late 2^nd^ FL, early 3^rd^ FL, late 2^nd^ FBM, early 3^rd^ FBM and ABM. In total, 247,565 cells among 285,641 cells passed the quality control and doublet removal **(Figures 3A and S5A)**. To reproduce the frequency of each cell type in pHSPC, we down-sized sample cell numbers analyzed based on the pHSC/MPP vs pHPC frequency from our FACS data (**Figure 3A and Table S2**).

**Figure 3;.**
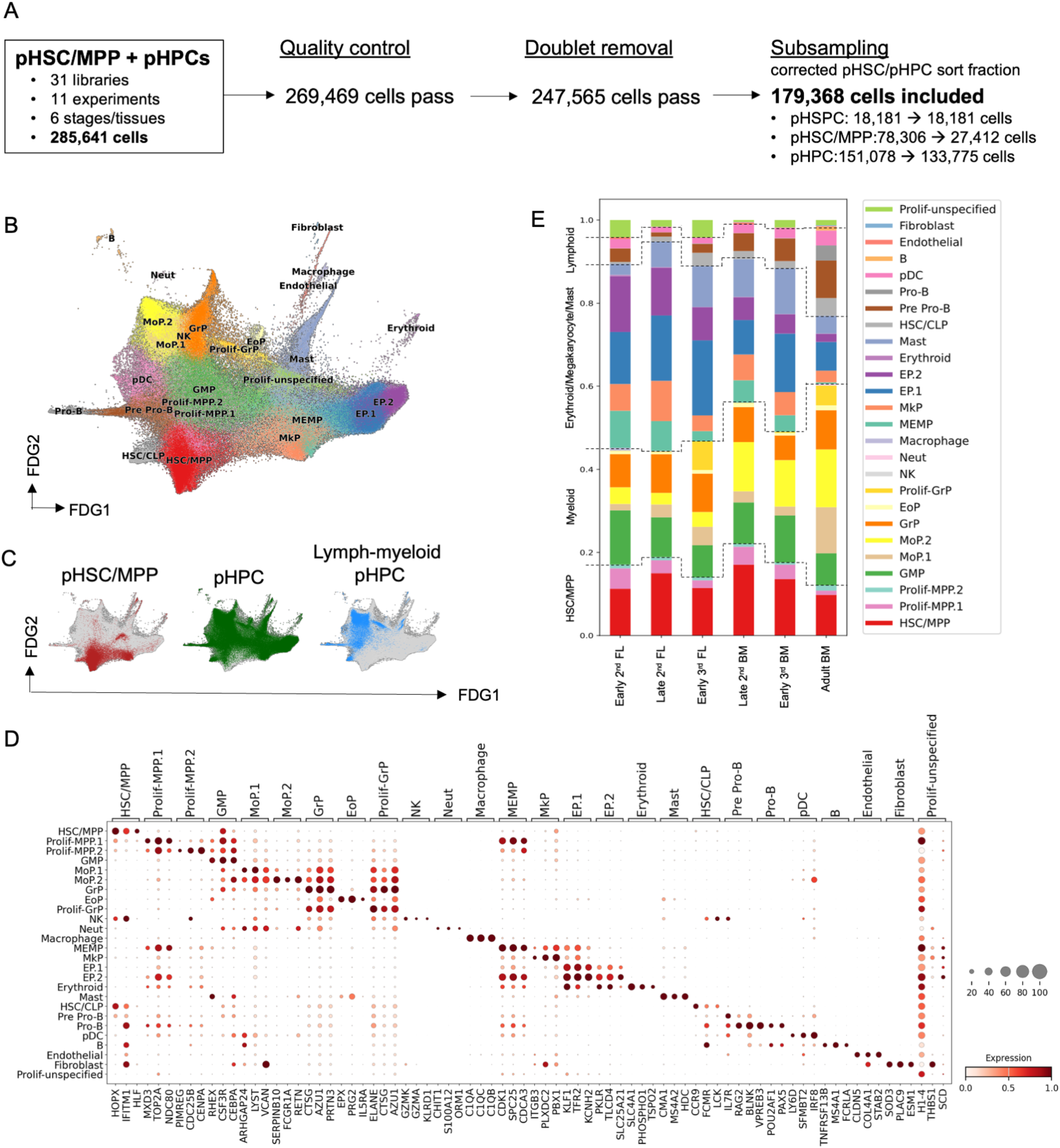
Transcriptional landscape of CD34^hi^ hematopoietic cells. A: Pipeline of the data analysis of 10x Genomics scRNA-seq. B: Force-directed graph (FDG) visualization of integrated CD34^hi^ cells, which is composed of pHSC/MPP and pHPC from multiple developmental stages/tissues; early 2^nd^-FL, late 2^nd^-FL, early 3^rd^-FL, late 2^nd^-FBM, early 3^rd^-FBM and ABM. Colors indicate cell types; Prolif-MPP, proliferating-MPP; MoP, monocyte-progenitor; GrP, granulocyte-progenitor; Neut, neutrophil; Prolif-GrP, proliferating-granulocyte-progenitor; EoP, eosinophil-progenitor; MkP, megakaryocyte-progenitor; EP, erythroid-progenitor. C: Distribution of pHSC/MPP, pHPC and lymph-myeloid pHPC cells on FDG. D: Gene expression of top 3 ranked genes of Wilcoxon-rank-sum test for each cell type. Dot color indicates log-transformed, min-max normalized gene expression value. Dot size indicates the percentage of cells in each category expressing a given gene. E: Cell type composition by developmental stage/tissue as the mean percentage of each cell type per each stage/tissue. Colors indicate cell types as shown in B.

This led to a final number of 77,017 early 2^nd^ FL cells (n = 2), 24,785 late 2^nd^ FL cells (n=2), 4,512 early 3^rd^ FL cells (n=1), 27,700 late 2^nd^ FBM cells (n=2), 17,935 early 3^rd^ FBM cells (n=1) and 27,419 ABM cells (n=4) that were included for downstream analysis. UMAP visualization integrating all these samples without batch correction revealed a batch effect across 10X library-preparation experiments. However, the batch effect was minimal across 10X library prep kit or sample donors if they are processed within the same experimental day (**Figures S5B, S5C, S5D and S5E**). Therefore, to minimize technical batch effects, while preserving the biological variation from the various developmental stages and FACS-isolated populations, the data were integrated using scvi-tools^39^ and visualized using dimensionality reduction methods of UMAP and Force-Directed Graph (FDG) (**Figures 3B**). According to the batch-corrected data integration, we performed Leiden clustering and identified differentially expressed genes (DEGs) using Wilcoxon rank sum test with Bonferroni correction to annotate cell clusters. The clusters were manually annotated by the expression pattern of previously characterized marker genes in human and NHPs, and 27 clusters of specific cell types including stem-progenitor-mature blood lineages were identified (**Figures 3B, S5F, S5G and Table S3**)^8,9,11,12,40^. Of note, we did not detect any differences of cell distribution when comparing between female vs male **(Figure S5H),** while we detected the apparent decrease in the frequency of granulocyte progenitor (GrP) population in thawed samples compared to fresh samples. **(Figure S5I)**. This is consistent with the result that CFU-GM recovery rate was decreased after thawing although there are few reports comparing the recovery rate of HSPC after freeze-thaw manipulation^41^.

The HSC/MPP cluster showed high expression of HOPX, IFIT1, and HLF, while the HSC/CLP cluster was distinct, expressing both HSC marker genes and lymphoid lineage related genes such as IL7R, LCK or LTB. We also found distinct myeloid progenitors, neutrophils, monocytes, macrophages, and plasmacytoid dendritic cells (pDCs) clusters. The GMP cluster expressed myeloid-lineage-specific genes (RHEX, CSF3R, and CEBPA) and GrP clusters expressed CTSG and AZU1. Monocyte progenitor (MoP) 1 and 2 clusters expressed SERPINB10, FCGR1A, or ZEB2 and erythroid progenitor (EP) 1 and 2 clusters expressed erythroid-related genes (KLF1, TFR2, PKLR, or TLCD4). The megakaryocyte progenitor (MkP) cluster expressed megakaryocyte-related genes (ITGB3, PLXDC2, and MMRN1), while the recently identified megakaryocyte-erythroid-mast cell progenitor (MEMP)^8^, share some genes with erythroid, megakaryocyte and mast cell lineages. The pre pro-B cell cluster were expressing IL7R, EBF1, and RAG2 and the pro-B cell cluster had an increasing expression of the B cell transcripts, CD19 and CD79B **(Figures 3D, S5G, Tables S4)**. Importantly, pHSC/MPP cells overlapped mostly in annotation with HSC/MPP (62.4 %), Prolif-MPP.1/2 (10.4 %) and MkP (9.5%) clusters. Likewise, pHPC cells were distributed across lineage-committed progenitors. Especially, CD45RA^+^CD90^-/mid^ fraction of pHPC, which was named lymphoid-myeloid HPC, was actually distributed on lymphoid and myeloid progenitor clusters **(Figure 3C)**. We also confirmed that surface marker genes used for HSPC purification **(Figure 2D)** were highly expressed in each designated stem or progenitor cell clusters as expected **(Figure S5G)**.

From our analysis, we also observed that the frequency of HSC/MPPs and each progenitor cluster dynamically shifts with developmental stage (**Figures 3E and S5J**). The frequency of the HSC/MPP cluster peaked at late 2^nd^ trimester for both FL and FBM, which is also consistent with our flow cytometry data (**Figure 2B**). The frequency of myeloid and lymphoid progenitors increased with age, while erythroid and megakaryocyte progenitors decreased with age. Interestingly, MEMP decreased with developmental stage and almost disappeared in ABM (**Figures 3E**) although MEMP has not been compared between FL and ABM in human.

Thus, we have successfully established the transcriptome of cynomolgus HSPCs, that has revealed the transcriptomic heterogeneity within HSPCs, which is in good agreement with recent studies showing the heterogeneity of human FL, FBM and ABM hematopoietic cells^8,9,10,11,12^.

### HSCs/MPPs can differentiate directly to MkPs

Next, we inferred lineage trajectories of cynomolgus hematopoiesis based on force directed graph (FDG) with partition-based approximate graph abstraction (PAGA). This showed the HSC/MPP cluster as the tip of the trajectory, with mainly three distinct trajectories downstream of HSC/MPP cluster; lymphoid, myeloid and megakaryocyte-erythroid-mast pathways **(Figures 4A and 4B)**. Interestingly, we also discovered a direct pathway from HSC/MPP to MkP cluster **(Figure 4C)** in addition to these three pathways passing through the proliferative state of progenitors, named Prolif-MPP.1/2. To investigate the molecular signatures along differentiation of HSC/MPP-MkP, HSC/MPP-MEMP-MkP, HSC/MPP-MEMP-EP and HSC/MPP-MEMP-mast pathways, we performed pseudotime analysis (Monocle3) to determine genes dynamically changing across pseudotime inferred differentiation pathways (**Figures 4D and S6, and Tables S5, S6, S7, S8**). Pseudotime analysis revealed similarities in HSPC differentiation to human hematopoiesis, and also further supported the existence of a previously uncharacterized direct differentiation pathway from HSC/MPP to MkP, that is distinct from the classical HSC/MPP-MEMP-MkP pathway.

**Figure 4;.**
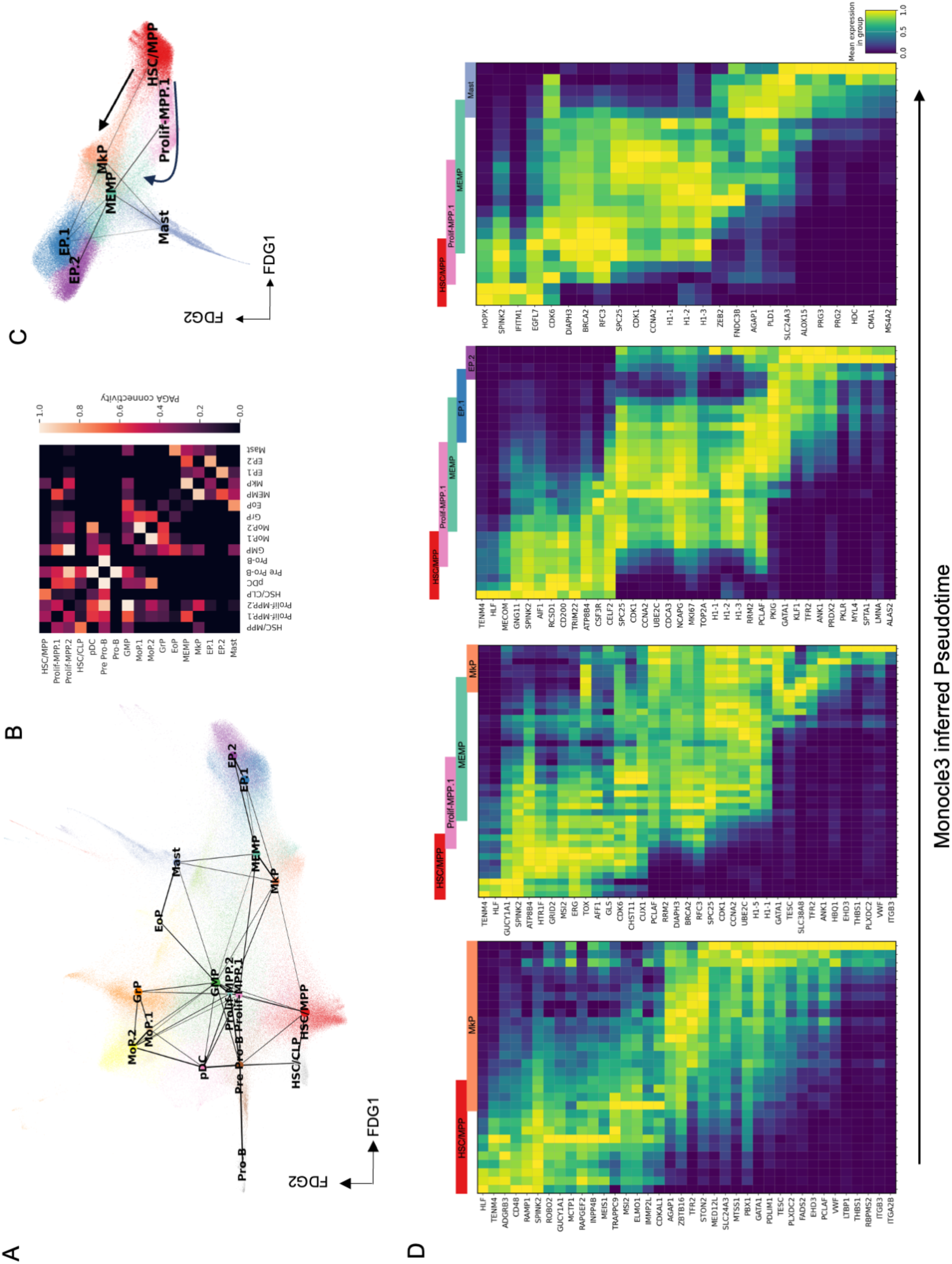
Differentiation trajectory of cynomolgus hematopoiesis. A: PAGA trajectory model imposed on the FDG visualization of the differentiation trajectory. Lines show connections; line thickness corresponds to the level of connectivity (low (thin) to high (thick) PAGA connectivity). Colors indicate cell types as shown in Figure 3B. B: PAGA connectivity scores of the populations shown in Figure 4A. C: FDG visualization of HSC/MPP, Prolif-MPP.1, MEMP, MkP, Mast, EP.1 and EP.2 from Figure 4A. D: Heat map showing min - max normalized expression of statistically significant (p < 0.001), dynamically variable genes from monocle3 inferred pseudotime analysis for MkP indirect differentiation pathway, MkP direct differentiation pathway, EP differentiation pathway and mast cell differentiation pathway.

### Transcriptional landscape of MPP across developmental stages and tissues

To establish the precise hierarchy of cynomolgus hematopoiesis, we reanalyzed all pHSC/MPP cells from 16 libraries/11 experiments (**Figure 5A**). In total, 87,543 cells passed quality control and doublet removal, with the remaining cells consisting of 46,768 early 2^nd^ FL cells (n=6), 10,867 late 2^nd^ FL cells (n=2), 849 early 3^rd^ FL cells (n=1), 8,649 late 2^nd^ FBM cells (n=2), 12,490 early 3^rd^ FBM cells (n=1), and 7,920 ABM cells (n=4). We identified 20 distinct cell clusters (**Figure 5B**), whereby cell types were annotated based on transferred cell labels from previous analysis **(Figures 3B and S7A),** and DEGs calculated by Wilcoxon rank sum test with Bonferroni correction **(Figure S7B, Tables S9 and S10)**. We found 6 clusters which originally reside within the HSC/MPP cluster, and we annotated them as HSC and MPP.1-5 **(Figures S7C and S7D).** We found that CD90 and CD31 gene expression levels were higher in the HSC and MPP.3 cluster than the others, and CD33 was not expressed in HSC and MPP clusters **(Figure S7E)**. Trajectory analysis revealed the HSC cluster as the top of the differentiation trajectory and strongly connecting to MPP clusters **(Figures 5C and 5D)**.

**Figure 5;.**
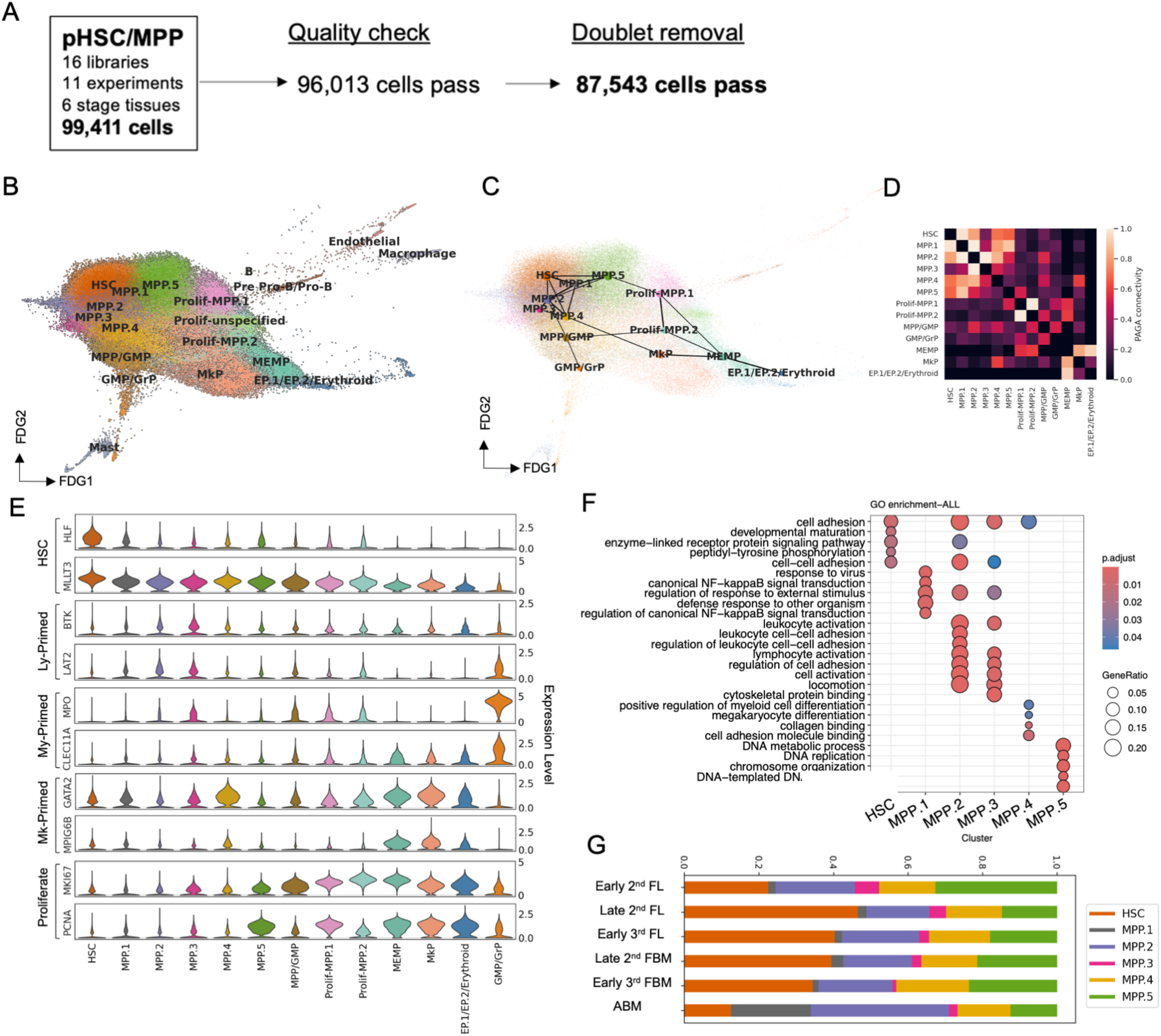
Transcriptional heterogeneity of pHSC/MPP. A: Pipeline of data analysis. B: FDG visualization of pHSC/MPP derived samples of multiple developmental stage/tissues. C: PAGA trajectory model imposed on the FDG visualization shown in Figure 5A. D: PAGA connectivity scores of the populations shown in Figure 5C. E: Violin plots showing ln-normalized gene expression of selected genes in each cell type. Colors correspond to those in Figure 5B. F: HSC and MPP clusters composition by developmental stage/tissue as the mean percentage of each subtype per each stage/tissue. G: Gene ontology analysis of genes upregulated across HSC and MPP clusters (> 0.5 log-fold change, q value < 0.05, fraction of cells within each group > 0.01) of pseudobulk populations from HSC and MPP clusters.

HSC signature genes such as HLF and MLLT3 were highly expressed in the HSC cluster and their expression levels decreased with differentiation into MPPs and oligopotent progenitors. Interestingly, among MPP clusters, MPP2/3 tend to show relatively increased expression of lymphoid-related genes such as SELL, BTK and LAT2, while MPP.3 concomitantly expressed myeloid-related genes such as MPO and CLEC11A. MPP4 showed increased expression of megakaryocyte-related genes such as GATA2 and MPIG6B (**Figure 5E).** Of note, all MPP clusters expressed HSC signature genes and MPP.1/2/3/4 remained in G1 phase **(Figure S7F).** This suggests that lineage priming may occur at the stage of non-cycling MPP. Moreover, a specific subset of MPP seemed to possess different lineage-priming programs simultaneously, and lymphoid-lineage priming seemed to start relatively earlier than myeloid-lineage priming. Gene Ontology (GO) enrichment analysis of HSC and MPP1/2/3/4/5 clusters also revealed that MPP2 is primed to leukocyte and lymphocyte and that MPP4 is a myeloid-primed lineage (especially megakaryocyte-primed lineage) (**Figure 5F, Tables S11 and S12**).

The frequency of HSC cluster peaked at late 2^nd^ trimester and decreased gradually from late 2^nd^ trimester to adult **(Figure 5G)**. This pattern is consistent with the frequency of pHSC (**Figure 2B)**. Interestingly, the MPP1 cluster is detected only in ABM. The significantly enriched GO terms indicate that MPP1 is related to canonical NF-kB and response to infection, which likely reflects the chronic exposure to various infections over the years and aging.

Taken together, we show that pHSC/MPP are heterogeneous, with some MPP clusters being already primed to certain lineages. The MPP cluster primed to megakaryocyte lineages connects the HSC cluster and the MkP cluster, supporting our data of the direct differentiation pathway from HSC/MPP to MkP and of a recent report claiming transcriptional lineage priming in fetal MPPs^8^.

### Characterization of transcriptionally-purified HSCs

Having established transcriptionally-purified HSCs from our transcriptome analysis **(Figure 5B and Table S13**), we then investigated the specific gene signatures of HSCs across developmental stages and tissues. Although most gene expression levels were similar across stages, some genes displayed distinct stage specific molecular profiles **(Figure 6A)**. We observed that TMSB10 and LDHA were increased in early 2^nd^ FL HSC cluster (**Figure 6B**). TMSB10 is involved in embryonic blood vessel formation and cell migration in various type of cells ^42,43^, which may reflect on early stages of HSC maturation and migration. LDHA is a direct target gene of HIF-α which activates erythropoietin or vascular endothelial growth in response to hypoxia^44^. Remarkably, GO terms related to mitochondria are enriched in early 2^nd^ FL-HSCs (**Figure 6C and Table S14**), which is consistent with several reports showing that FL-HSCs contain more mitochondria with higher bioenergetic function than adult HSCs and are more dependent mitochondrial metabolism^45^. FOS, JUND and DUSP1were highly expressed at late 2^nd^ FL and ABM, and these gene are well-known to be involved in cell proliferation, myeloid differentiation or aging^46,47,48^ (**Figure 6B**). We also detected several genes that were increased in ABM; AR, which is related with hematopoietic differentiation and ANXA1, which is an anti-inflammatory molecule (**Figure 6B**). The significantly enriched GO terms indicated that ABM-HSC is related to canonical NF-kB signal transduction and regulation of canonical NF-kB signal transduction (**Figures 6C and Table S13**), probably due to increased NF-kB activity during aging and inflammation^49^.

**Figure 6;.**
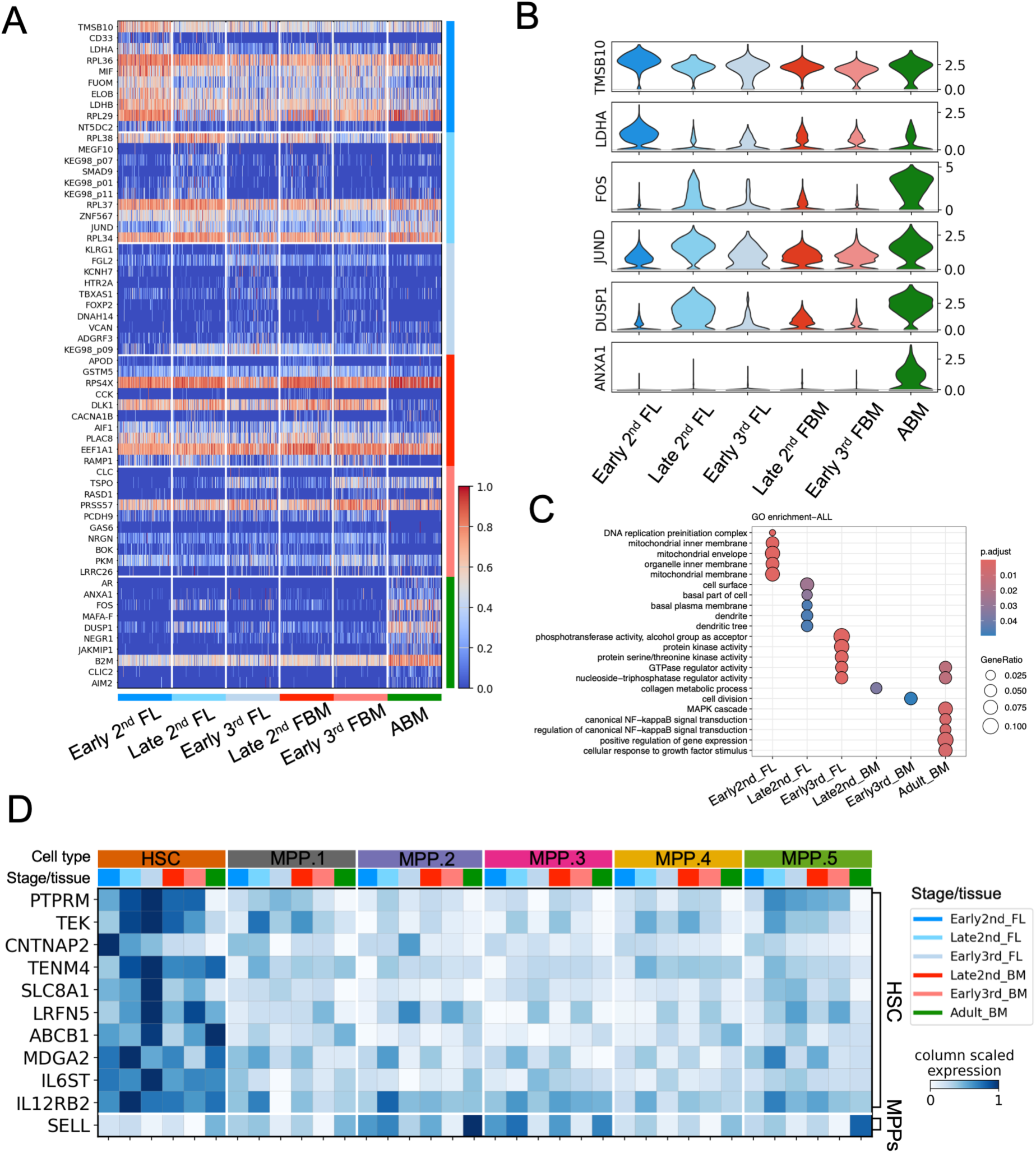
Characterization of transcriptionally-purified HSC. A: Heatmap showing top 10 ranked DEGs (> 0.5 log-fold change, q value < 0.05, fraction of cells within each group > 0.01) for each developmental stage/tissue when compared in transcriptionally-purified HSC. B: Violin plots showing ln-normalized gene expression of selected genes in each developmental stage/tissue. C: GO analysis of genes upregulated upon development (> 0.5 log-fold change, q value < 0.05, fraction of cells within each group > 0.01) of pseudobulk populations from each stage/tissue in transcriptionally-purified HSC. D: Matrix plot showing significantly differentially expressing surface marker genes. Matrix color indicates log-transformed, min-max normalized gene expression value.

Finally, for further purification of HSCs, surface proteins are necessary. Therefore, we investigated genes coding surface proteins that were specific to the HSC cluster compared to the MPP clusters. By comparing HSCs vs MPP.1-5 by Wald tests, we detected 10 surface marker genes for purifying transcriptionally-purified HSC and SELL for remove MPP clusters (**Figure 6D**) across the developmental stages from fetus to adult. In conclusion, we have successfully isolated multiple, previously uncharacterized HSC markers that can potentially be used for flow cytometry and further analysis of cynomolgus HSCs.

## Discussion

In this study, we have extensively characterized hematopoiesis and HSC development in the cynomolgus monkey, focusing predominantly on HSCs/MPPs, to establish a comprehensive HSC dataset that covers gestational periods of the NHP, previously not analyzed in human. We functionally demonstrate, through HSC xenotransplant assays, the existence of HSCs in the FL and FBM during specific gestational stages, recapitulating events seen in human. Furthermore, by establishing a defined gating strategy (Lin^-^CD45^+^CD34^hi^CD45RA^-^CD33^-^CD31^hi^CD90^hi^), we successfully purified transplantable HSCs and generated the transcriptome of HSPCs, describing a previous uncharacterized direct differentiation pathway from HSC/MPP to MkP. Furthermore, we were also able to distinguish the distinct molecular characteristics of highly-purified HSCs and lineage-primed MPPs, through which we could identify potential uncharacterized candidate surface markers for further HSCs purification.

Understanding the principal sites of hematopoiesis during development, including its duration and characteristic properties, is crucial when consider cynomolgus monkeys as a preclinical model to investigate processes such as fetal-maternal immune or metabolic response^50,51^. In human, FL hematopoiesis is predominant until mid-2^nd^ trimester and FBM hematopoiesis becomes dominant after 20 post-conception weeks (PCW)^8,52^, which corresponds approximately to E86-E90 in cynomolgus monkeys. Indeed, similar to humans, we observed that cynomolgus HSCs shift from FL to FBM during mid-late 2^nd^ trimester. Surprisingly, we found that early 3^rd^ trimester E121 FL still have repopulation capacity, which was longer than expected^53^. Hence, due to the accessibility of samples at later stage of pregnancy in cynomolgus monkeys, we have successfully characterized fetal hematopoiesis —especially at later stages that are currently not possible in human^8^, — potentially shedding light on events occurring in human and further extending our understanding of human fetal development.

We identified similarities and differences in HSC surface markers expressed between cynomolgus monkeys and humans. Perhaps, the most significant difference we found was that human HSCs are negative for CD38, whereas cynomolgus HSCs are positive for CD38. This is critical because previous studies have used CD38 negativity as a HSC marker in NHPs^54,55^. Human CD33 is expressed on both HSCs and AML cells and thus gene editing of CD33 on HSCs is expected to have protective effects on HSCs from anti-CD33 AML therapies such as CAR-T^56^ and rhesus macaque have also been used as a preclinical model for this research purpose^57,58^. However, as our results point out the critical limitations of using cynomolgus monkeys for such research purposes, this suggests that better characterization of CD33 marker expression in HSCs is necessary for preclinical studies using NHPs.

Historically, in vitro colony formation and transplantation assays using flow cytometric antibodies were performed for the identification of markers of HSCs. When necessary, antibodies were then produced based on gene expression data, leading to the further purification of HSCs. Regarding transplantation assays, allogeneic transplantations are typically performed in mice, whereas in humans and NHPs, xenotransplantation is the gold standard— though the differences in recipient immune system and niche environment has presented challenges in accurately assessing multi-lineage and long-term engraftment. In our study, we achieved the identification and functional validation of transplantable HSCs in the xenotransplantation setting. Therefore, the next step will be to move to an autologous setting and evaluating the repopulation potential of cynomolgus HSCs. Furthermore, we anticipate that the 10 candidate surface markers identified from our study will become useful for purifying cynomolgus HSCs in the future since there are limited commercially available antibodies that can be currently used in cynomolgus monkeys for flow cytometry.

Our extensive analysis on CD34^hi^ HSPCs revealed divergent cells within the HSPC population and we observed a dynamic shift of their compartments across developmental stages and tissues. Furthermore, we found that HSCs/MPPs and MkPs were directly connected based on FDG and PAGA analysis, suggesting the possibility of a direct differentiation pathway from HSC/MPPs to MkPs without passing through any intermediate MEMP step. This is consistent with myeloid-bypass pathway in mice^59^ and direct differentiation of Mk from HSC and multipotency of MkP in humans^60^.

We also found that some MPPs are already primed to specific lineages. Interestingly, our analysis on FDG and PAGA suggest that the primary bifurcation occurs at the MPP stage into Mk-primed MPP (MPP.4) or lympho-myeloid MPP (MPP.2) as suggested in human fetal hematopoiesis^60^. The existence of MPP1 may reflect that HSCs in ABM produce MPPs using canonical NF-kB activation and/or under some infection and/or inflammation events such as aging.

In summary, we anticipate that our comprehensive dataset will serve as a basis for building an HSC atlas of cynomolgus monkeys that will facilitate a better understanding of conserved and non-conserved properties as well as mechanisms between NHPs and humans, which will be necessary for better translational applications in future.

### Limitations

In this study, many assays and experiments were performed on early 2^nd^ FL, and we could not increase the number sampling time points during embryonic developmental stages or increase donor numbers on later stages of development because of the maternal physical burden associated with fetus sampling and financial cost. Sampling of earlier stages also requires hysterectomy leading to maternal euthanasia and increases abortion risk in later stages. For adult bone marrow sampling, we used female adult cynomolgus macaque that were planned to undergo euthanasia for other research purposes and thus the sampling time point was restricted to 9-13-year-old.

As we focused on more on immature populations such as HSCs/MPPs, functional analysis was limited to transplantation assays, but if we were to characterize the differentiation potential of oligo-mono potent progenitors or more mature blood cells, this would require further experiments such as in vitro colony assays.

## Supporting information

Key resource table

## Acknowledgements

We thank S. Goulas for critical reading and constructive suggestions on our manuscript; members of the Single-cell Genome Information Analysis Core (SignAC) at WPI-ASHBi for their support. RY was supported by SUNTORY SunRise.

## Author contribution

S.O. and R.Y. conceived and designed the project. S.O. contributed to the performance of the experiments and analysis of the data. S.O. and R.S. performed bioinformatics analysis. S.O. and K.T. performed scRNA-seq library preparation and sequencing. K.M., T.N., M.N. and T.T. provided cynomolgus macaque samples. I.K., C.I. and H.T. performed cynomolgus macaque breeding, embryo manipulation and cesarian section. S.B., A.T-K., M.S. and Y.M. advice and interpreted data. S.O. and R.Y. wrote and edited the manuscript. R.Y. supervised the study. All the authors reviewed and edited the final version of the manuscript

## Methods

### Animals

All animal experiments were performed with the ethical approvals of Kyoto University and Animal Care and Use Committee of Shiga University of Medical Science.

For the animal experiments using fetus cynomolgus monkeys, the manipulation of cynomolgus embryo, pregnancy check and cesarean section was performed as described previously^61,62^. The sex of fetus was confirmed by checking the gross appearance of its gonads or by PCR for UTX/UTY gene^62^. In this study, 1 late 1^st^ trimester fetus, 31 early 2^nd^ trimester fetus, 2 late 2^nd^ trimester fetus, 2 early 3^rd^ trimester fetus and 11 adults were used **(Table S15)**.

We collected ABM samples from female cynomolgus macaques scheduled for a euthanasia for another research purpose. The monkeys were anesthetized with an intramuscular injection of ketamine hydrochloride (5 mg/kg) and xylazine hydrochloride (1mg/kg). Post-hysterectomy, we performed bone marrow aspirations from bilateral iliac crests, limiting the volume to a maximum of 30ml to avoid impacting respiratory and circulatory dynamics. The procedure was followed with euthanasia by exsanguination.

Immunodeficient NOG-W41 (formal name; NOD.Cg-*Prkdc^scid^ Il2rg^tm1Sug^ Kit^em^*^1^(V831M)*^Jic^*/Jic) and NSG (formal name; NOD.Cg-*Prkdc^scid^ Il2rg^tm1Wjl^*/SzJ) mice were purchased from Central Institute for Experimental Animals and Jaxson Laboratory, respectively. In all experiments, similar number of recipient male and female mice were used to avoid potential bias.

Cynomolgus macaque were housed individually in appropriate cages under a 12-h light/12-h dark cycle. Temperature and humidity in the animal rooms were maintained at 25±2°C and 50±5%, respectively. Mice were housed in a specific pathogen-free animal facility under a 14-h light/10-h dark cycle.

### Tissue processing and cell isolation from FL and FBM

All tissues were processed immediately after isolation. FL tissues were cut into smaller fragments, placed onto 70µm cell strainers, and mechanically dissociated using a 2.5mL syringe plunger, followed by a rinse with ice-cold PBS. For FBM collection, bilateral femurs, tibias, and humeri were cleaned of soft tissues, placed in a ceramic mortar containing ice-cold PBS, cut into smaller fragments, and manually crushed with a ceramic pestle. The crushed samples were then transferred to 70µm cell strainers to filter out bone fragments and rinsed with ice-cold PBS. Filtered cells from dissociated FL and FBM were collected by centrifugation at 400g for 5 minutes at 4°C.

### Mononuclear cell isolation from ABM

ABM, aspirated with 5000u/ml heparin, was diluted 2-fold with room temperature PBS and transferred to a 70mm cell strainer to filter out bone fragments and clogs. By using a 95% diluted lymphoprep gradient, mononuclear cell layer was collected after centrifugation at 1500g for 30 minutes at room temperature and washed with ice-cold PBS.

### CD34 enrichment with MACS separation system

FL, FBM and ABM cells were resuspended in ice-cold PBS with 5ul of Fc receptor blocker (Human TruStain FcX, Biolegend) and were incubated for 10 min at 4°C. CD34^+^ cells were labeled with APC or APC-Cy7 conjugated anti-CD34 antibodies (clone 561; BD Pharmingen, San Jose, CA, USA) at a concentration of 1:20 dilution and anti-APC MicroBeads at a concentration of 1ul/1×10^7^ cells. Labeled CD34^+^ cells were isolated using MACS separation system (Miltenyi Biotec, Bergisch Gladbach, Germany).

### Flow cytometry and cell sorting

Freshly isolated or frozen-thawed MACS-enriched CD34^+^ cells were stained with antibodies which were validated for binding to cynomolgus protein^32,51,63^. Cells were resuspended in 100 μl of PBS with antibody mix. Cells were stained for 30 min at 4°C, washed with PBS and resuspended at 1.0 × 10^7^ cells per ml. DAPI or PI was added to a final concentration of 1ug/mL immediately before flow cytometric analysis. Compensation setting was performed based on auto-compensation using beads following manufacture’s instruction (UltraComp eBeads Compensation Beads, Invitrogen). Flow sorting was performed on a BD FACSAria Fusion instrument using DIVA v.8.0.2 or 9.4, and data were analyzed using FlowJo (v.10.8.1, BD Biosciences). Cells were sorted into chilled 1.5ml microtubes. Each tube was prefilled with specific media: 200µl of sterile 0.1% PCL-PVAc-PEG (Soluplus)^64^ in PBS for xenotransplantation; 300µl of 10% fetal bovine serum (FBS) in IMDM for colony assays; and 500µl of 0.04% bovine serum albumin (BSA) in PBS for scRNA-seq, respectively.

### Mouse xenotransplantation in NOG-W41 or NSG mice and engraftment analysis

Xenotransplantation was performed on immunodeficient 5-13-week-old NSG mice with 2.5-2.8Gy irradiation or NOG-W41 mice without irradiation. NSG mice were used for identification of transplantable HSCs across multiple developmental stages and tissues (**Figure 1B**), and NOG-W41 mice were used unless otherwise mentioned. Freshly-isolated and/or enriched with MACS or cell sorting were injected from tail vein with 200-250 µl/body. Mice underwent retro-orbital bleeding monthly, and engraftments were identified by the presence of a distinct population of cynomolgus CD45^+^ cells, constituting more than 0.1% of total blood cells, three months post-transplantation. Severe combined immunodeficiency (SCID)-repopulating cells (SRC) frequency was calculated using the extreme limiting dilution analysis software (ELDA v1.5.0)^65^. For secondary transplants, a single-cell suspension of bone marrow cells from primary recipient mice was prepared, enriched for CD34^+^ cells using MACS, and then transplanted into secondary recipient mice.

### Colony-forming cell assay

For colony-forming cell assays, 100-1000 sorted cells within 300 µl of IMDM were seeded into 3.0 ml ColonyGEL 1402 (ReachBio) and plated onto 35 × 10 mm culture dish at a volume of 1.0 ml/dish. Hematopoietic colonies were scored and classified morphologically after 14 days; colony forming unit (CFU)-granulocyte (CFU-G), -macrophage (CFU-M), -granulocyte-macrophage (CFU-GM), -erythrocyte (CFU-E), -granulocyte-erythroid-macrophage-megakaryocyte (GEMM), and burst forming unit-erythrocyte (BFU-E).

### 10X genomics library preparation and sequencing

FACS isolated cells were filtered via 40 um cell strainer and collected with centrifugation 400g for 10 minutes at 4°C. Cells were loaded onto each channel of a Chromium chip followed by GEM generation with Chromium X or Chromium Controller (10X Genomics). Library preparation (10X Genomics, Chromium Next GEM Single Cell 5’ Reagent Kits v.2, Chromium Next GEM Single Cell 5’ HT Reagent Kits v.2) were performed following the manufacturer’s instructions, with PCR cycle for cDNA amplification were adjusted based on sorted cell counts. Libraries were sequenced using NovaSeq 6000 platform (Illumina) to achieve a minimum depth of 40,000 raw reads per cell. The libraries were sequenced using the following parameters: Read1: 151 cycles, i7: 10 cycles, i5: 10 cycles; Read2: 151 cycles to generate 151-bp paired-end reads.

### Read alignment, quantification and quality control of scRNA-seq data

Raw files were processed with the “mkfastq” command in Cell Ranger (v. 7.0.0) and mapped to the cynomolgus monkey reference genome MFA1912RKSv2 with “count” command. Downstream processing of aligned reads was performed using Scanpy^66^ (v.1.9.5) following the online tutorials(https://scanpy.readthedocs.io/en/stable/tutorials.html). The low-quality cells and dying cells (cells with fewer than 500 detected genes and total mitochondrial genes expression exceeded 1%) were excluded. Genes that were expressed in fewer than 3 cells were also excluded. Approximately 4,500 genes per cell were detected.

### Doublet detection

Doublets score was calculated using DoubletDetection^67^ (v.4.2) with the default setting. Detected doublets which had p-value threshold < 1e-16 and voter threshold > 0.5 were removed.

### Subsampling

In order to reflect the actual frequency of cell types when analyzing the integrated data from “CD34^low^ + pHSC/MPP + pHPC” (**Figure S4**) or “pHSC/MPP + pHPC” (**Figure 3**), cells from each population were subsampled based of their frequency within FACS gate. This subsampling step was performed after quality control and doublet detection, using the “scanpy.pp.subsample” command implemented in Scanpy. Only subsampled cells were used for downstream analysis.

### Data integration

Before the data integration, the data was normalization to 10,000 reads per cell, log scaled, and higly variable genes were detected with “min_mean=0.0125, Max_mean=3, min_disp=0.5” using Scanpy. Batch correction was performed using scvi-tools^39^ (v.1.0.4) with the “batch_key” set to each independent cell sorting experiments, “n_hidden=256, n_latent=80, n_layers =2”. The resulting latent representations were employed for the following neighborhood graph construction and dimensionality reduction.

### Dimensionality reduction, leiden clustering and cell type annotation

The integrated data were subjected to “scanpy.pp.neighbors” command to construct the nearest-neighbor graph with a parameter “use_rep=“X_scVI”, n_neighbors = 30, n_pcs = 60”. Dimensionality reduction methods included UMAP using “scanpy.tl.umap” (min_dist =0.5, spread =1.0 for **Figures 3 and S4**, min_dist=1.0, spread =1.5 for **Figure 5**), and FDG using “scanpy.tl.draw_graph” which used the fa2 (v.0.3.5) Python Package. For the leiden clustering, scanpy.tl.leiden with the resolution at 1.0 was used. Cell types were assigned to individual clusters by checking the DEGs calculated by Wilcoxon rank-sum test with Bonferroni correction and expression pattern of previously characterized marker genes in human and NHPs.

### Trajectory analysis and pseudotime analysis

PAGA using “scanpy.tl.paga” (model = v1.2) and FDG with paga initialization was applied to infer the differentiation trajectories. We removed Neutrophil, B cell, Erythroid, Endothelial, Macrophage, Fibroblast, Prolif-GrP and Prolif-unspecified clusters on **Figure 3B**, Mast cell, Pre Pro-B, B cell, Macrophage, Endothelial, Fibroblast and Prolif-unspecified clusters on **Figure 5B**, and recalculated the PAGA graph. Genes that vary across pseudotime were calculated using the Graph-autocorrelation analysis (“graph_test” with neighbor_graph= “principal_graph” command) in Monocle3 (v.1.3.1)^68^. The significant genes for each gene modules were ranked by effect size according to the morans_I value, and top genes are shown in the heat map **(Figure 4D)**.

### GO analysis and differential expression analysis

GO analysis, comparing HSC and MPP subtypes or developmental stage and tissues withing HSC cluster, was performed using ClusterProfiler (v.4.10.0)^69,70^ implemented in R. The DEGs were calculated by comparing each cluster by Wilcoxon rank-sum test with Bonferroni correction. DEGs were defined as log fold change > 0.5, adjusted p-value < 0.05 and fraction of cells within each group > 0.01. For detecting surface maker genes **(Figure 6D)**, DEGs were calculated by comparing HSC cluster vs remaining MPP.1-5 clusters by diffxpy (v.0.7.4) using Wald test. Surface protein list for human were obtained from BioMart database (https://www.ensembl.org/info/data/biomart/index.html), then surface marker genes were selected among DEGs.

## Resource Availability

### Material Availability

This study did not generate new unique reagents.

### Data and Code Availability

The scRNA-seq raw data have been deposited at Sequence Read Archive (SRA) : PRJNA1090143, and the processed datasets have been uploaded at Gene Expression Omnibus (GEO) : GSE262140. They are publicly available and can be downloaded as of the publication.

This paper does not report original code

All other relevant information required to reanalyze the data reported in this study is available from the lead contact upon request

