## Supplementary figures and images for "Atlas of Cynomolgus Macaque Hematopoiesis"

### Sup Fig 1

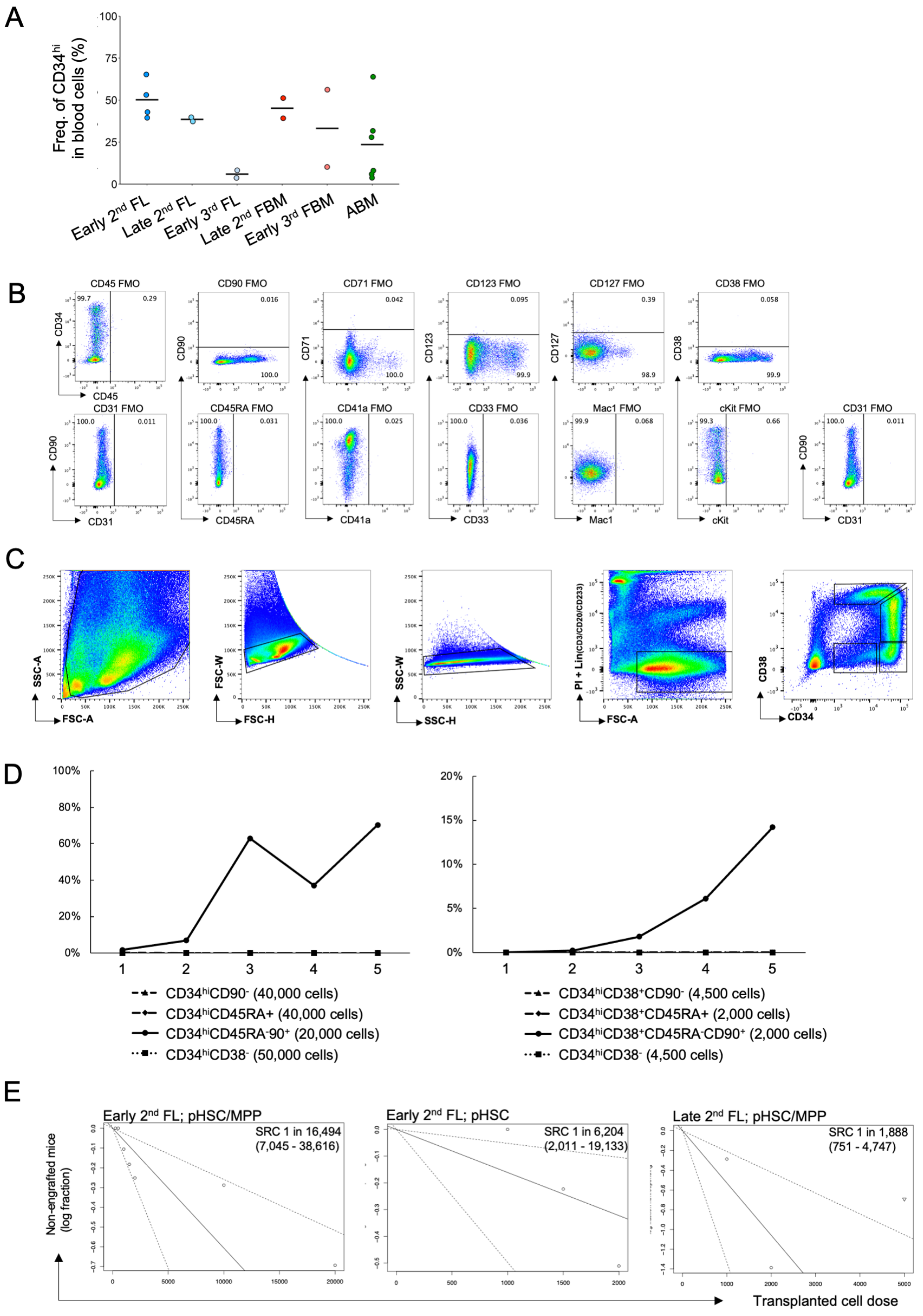

### Sup Fig 2

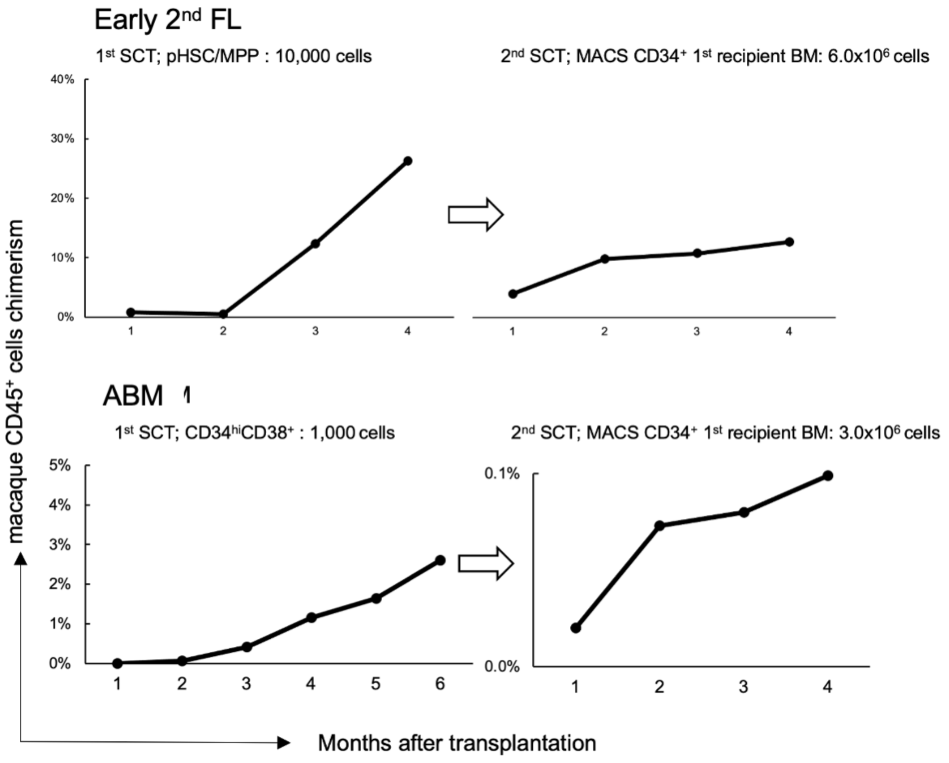

### Sup Fig 3

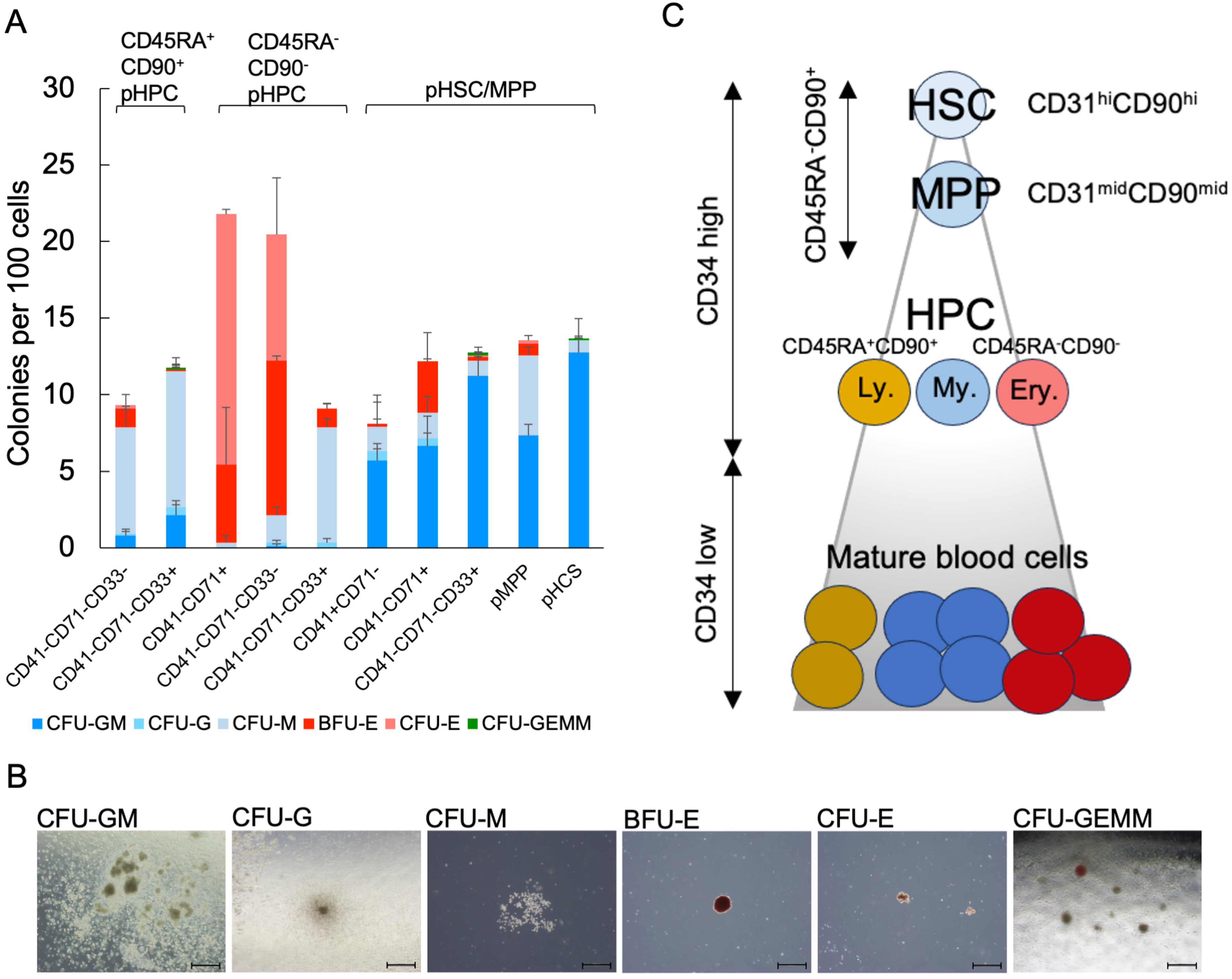

### Sup Fig 4

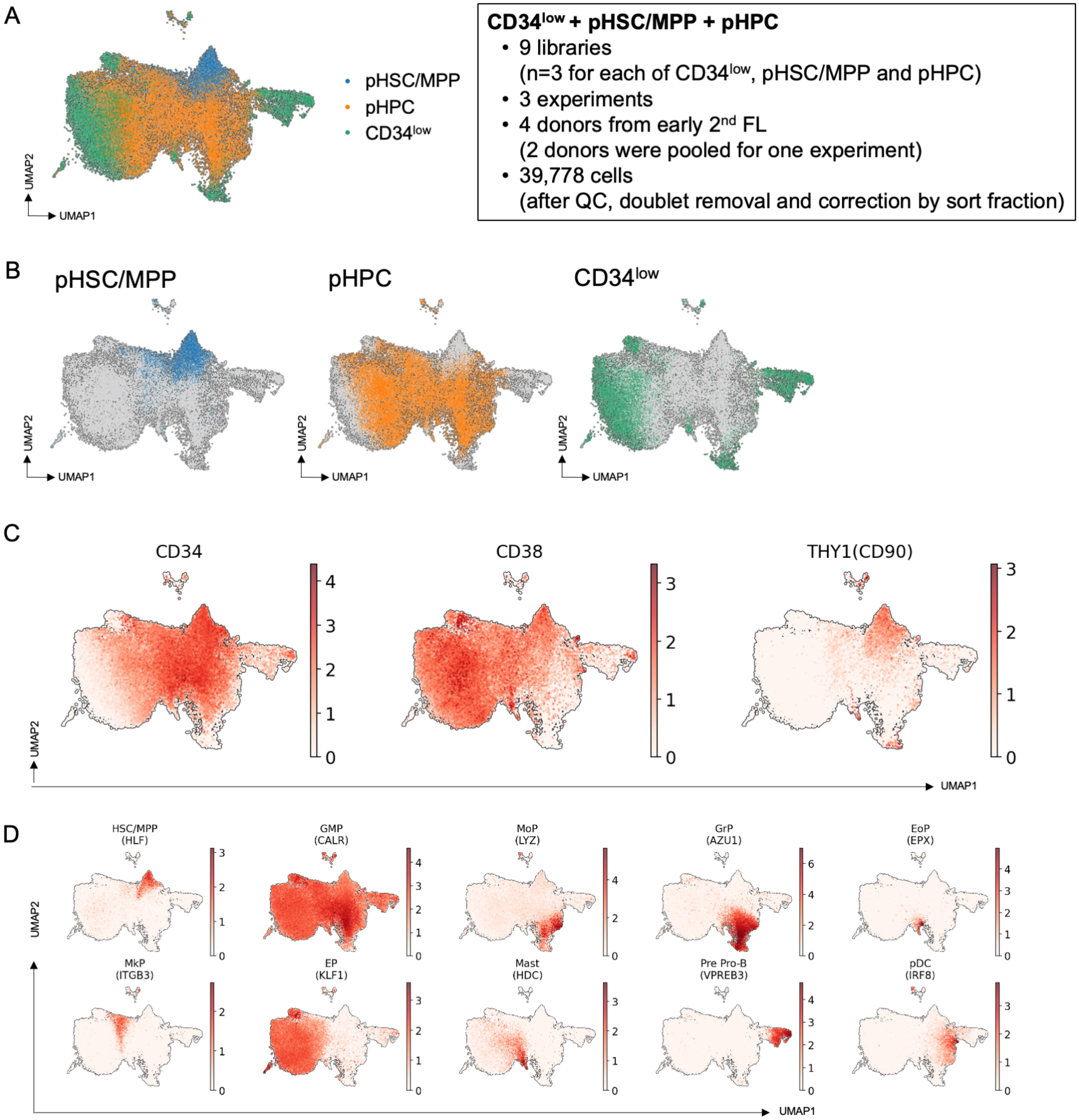

### Sup Fig 5

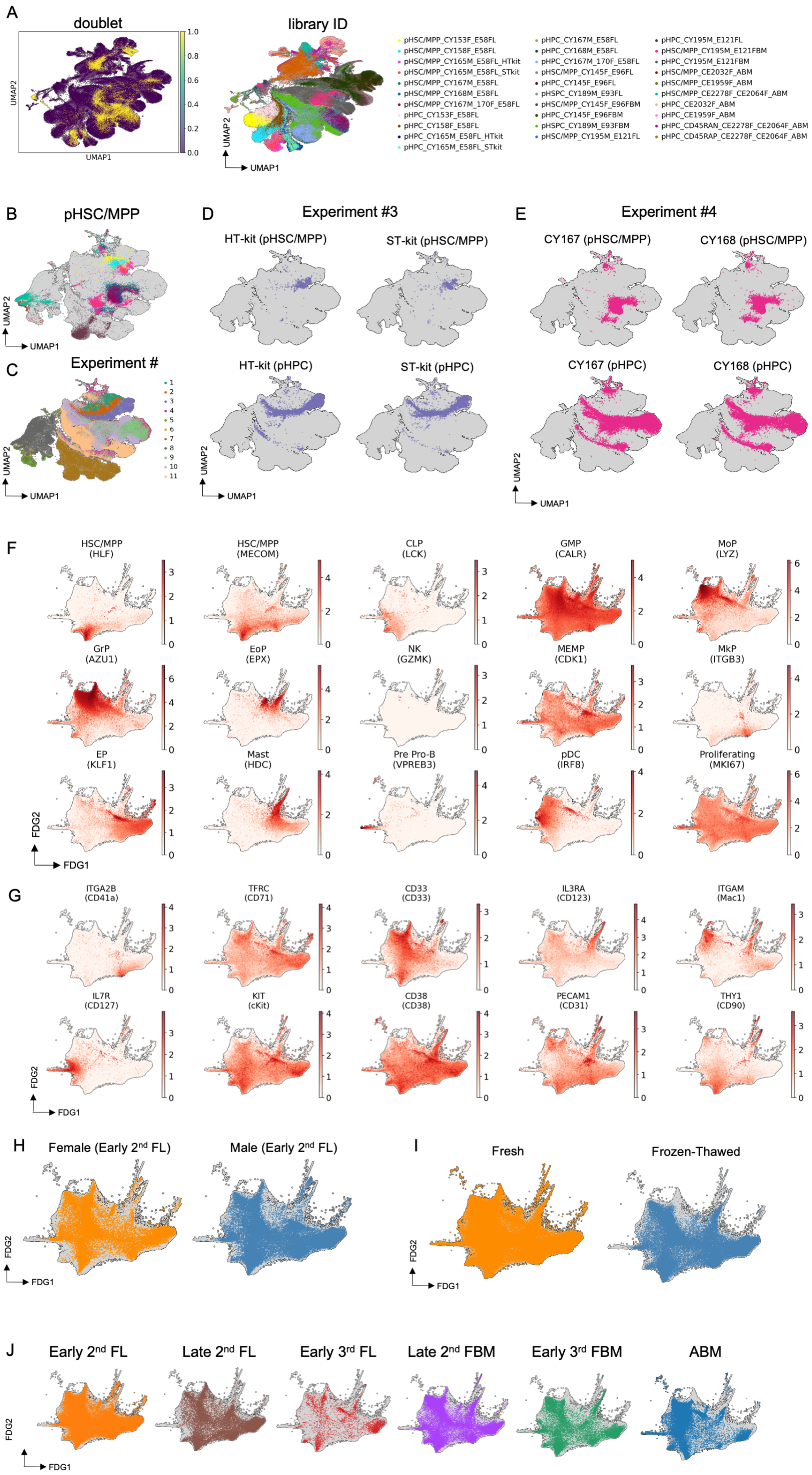

### Sup Fig 6

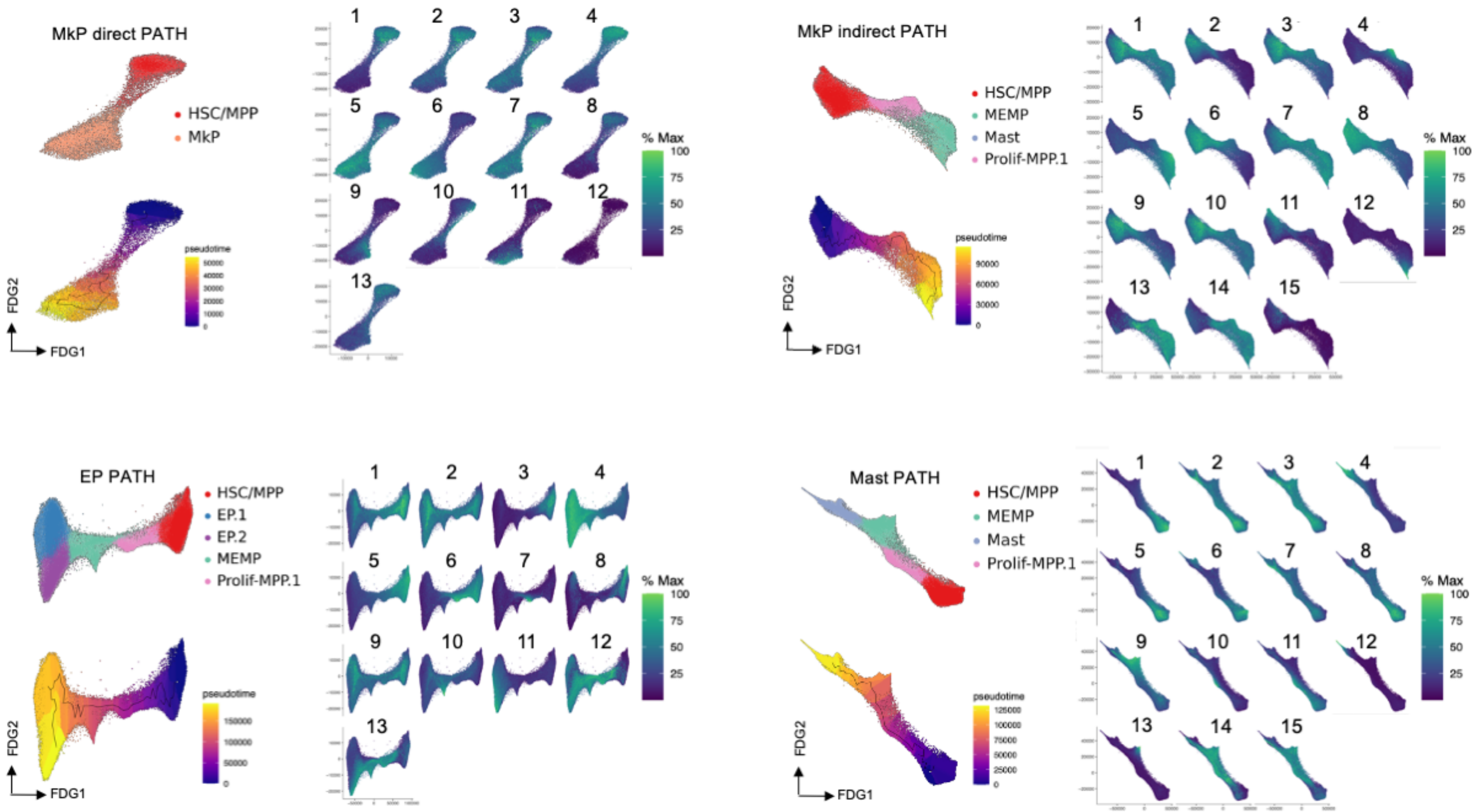

### Sup Fig 7

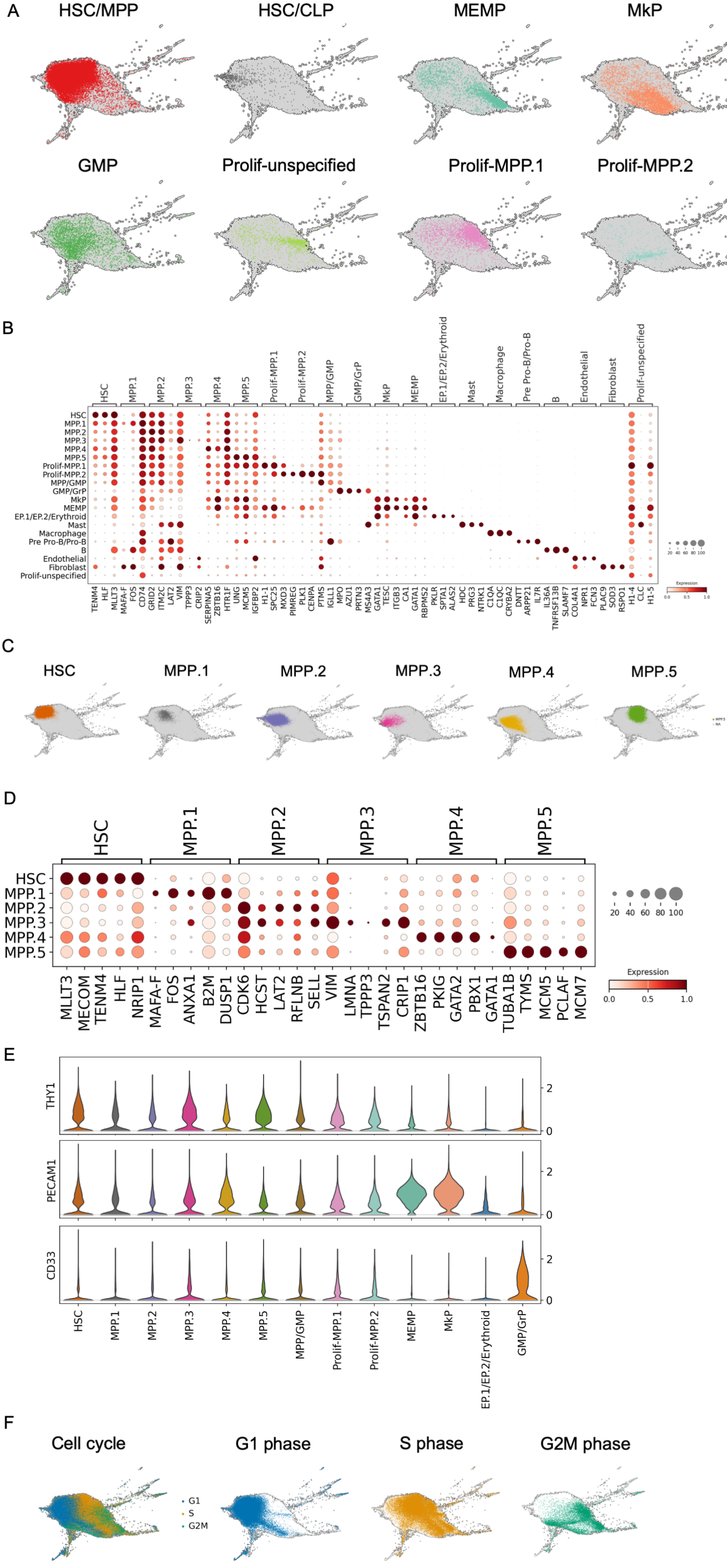

### Table 1

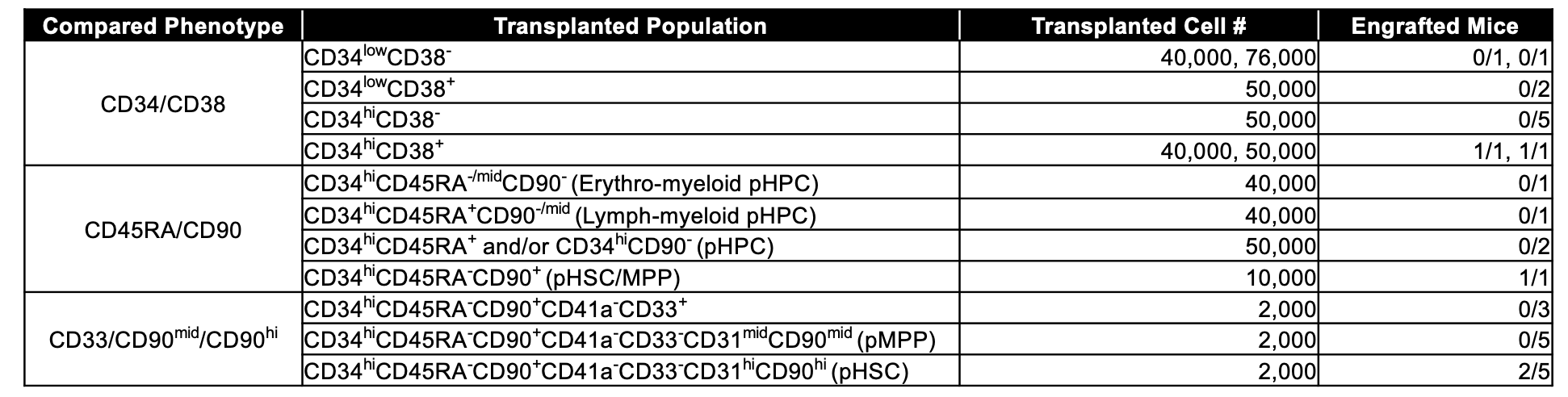
