## Supplementary material for "Atlas of Cynomolgus Macaque Hematopoiesis": Key resource table

| **REAGENT or RESOURCE** | **SOURCE** | **IDENTIFIER** |
| --- | --- | --- |
| **Antibodies** | | |
| FITC CD14(clone: M5E2) | BioLegend | Cat:301804, RRID:AB_314186 |
| FITC CD38(clone: AT-1) | Stemcell Technologies | Cat:60131FI, RRID:AB_2884030 |
| FITC HLA-ABC(clone: G46-2.6) | BD Biosciences | Cat:557348, RRID:AB_396654 |
| PE CD3(clone: 10D12) | Miltenyi | Cat:130-118-534, RRID:AB_2733803 |
| PE CD31(clone: WM59) | eBioscience | Cat:12-0319-42, RRID:AB_10669160 |
| PE CD33(clone: AC104.3E3) | Miltenyi | Cat:130-113-349, RRID:AB_2726124 |
| PE CD45(clone: D058-1283) | BD Biosciences | Cat:552833, RRID:AB_394483 |
| PE/Dazzle594 CD20(clone: 2H7) | BioLegend | Cat:302348, RRID:AB_2564387 |
| PE-Cy7 CD45RA(clone: 5H9) | BD Biosciences | Cat:561216, RRID:AB_10611721 |
| APC CD3(clone: SP34-2) | BD Biosciences | Cat:557597, RRID:AB_398626 |
| APC CD34(clone: 561) | BioLegend | Cat:343608, RRID:AB_2228972 |
| APC CD41a(clone: HIP8) | BioLegend | Cat:303710, RRID:AB_2249385 |
| APC CD90(clone: 5E10) | eBioscience | Cat:17-0909-42, RRID:AB_11042579 |
| AF700 CD11b(clone: ICRF44) | BioLegend | Cat:301356, RRID:AB_2750075 |
| APC-Cy7 CD31(clone: WM59) | BD Biosciences | Cat:563653, RRID:AB_2738350 |
| APC-Cy7 CD34(clone: 561) | BioLegend | Cat:343614, RRID:AB_2571927 |
| APC-Cy7 cKit(clone: 104D2) | eBioscience | Cat:47-1178-42, RRID:AB_2815166 |
| BV421 CD20(clone: 2H7) | BioLegend | Cat:302330, RRID:AB_10965543 |
| BV421 CD31(clone: WM59) | BioLegend | Cat:303124, RRID:AB_2563810 |
| BV421 CD42a (clone:ALMA.16) | BD | Cat:565444, RRID:AB_2739238 |
| eF450 CD123(clone: 7G3) | eBioscience | Cat:48-1238-42, RRID:AB_11151323 |
| BV510 CD41a(clone: HIP8) | BioLegend | Cat:303736, RRID:AB_2687213 |
| BV650 CD127(clone: A19D5) | BioLegend | Cat:351326,RRID:AB_2562095 |
| BV711 CD71(clone: L01.1) | BD Biosciences | Cat:745418, RRID:AB_2742970 |
| BV786 CD45(clone: D058-1283) | BD Biosciences | Cat:563861, RRID:AB_2738454 |
| BUV395 CD90(clone: 5E10) | BD Biosciences | Cat:563804, RRID:AB_2632398 |
| BUV737 CD123(clone: 7G3) | BD Biosciences | Cat:741778, RRID:AB_2871132 |
| Biotin CD14(clone: M5E2) | BioLegend | Cat:301826, RRID:AB_2291250 |
| Biotin CD20(clone: 2H7) | eBioscience | Cat:13-0209-82, RRID:AB_657690 |
| Biotin CD233(clone: BRIC6) | IBGRL | Cat: 9439BI |
| Biotin CD3(clone: 10D12) | Miltenyi | Cat:130-092-008, RRID:AB_871672 |
| **Chemicals, Peptides, and Recombinant Proteins** | | |
| 10%BSA | Sigma-Aldrich | A1595-50ML |
| STEM-CELLBanker EX GMP grade | Takara | CB091 |
| IMDM | Wako | 098-06465 |
| **Critical comercial assay** | | |
| ColonyGEL™ 1402, NHP Complete Medium | ReachBio | 1402 |
| Chromium Next GEM Single Cell 5' Kit v2, | 10x Genomics | PN-1000263 |
| Chromium Next GEM Single Cell 5' HT Kit v2 | 10x Genomics | PN-1000356 |
| High Sensitivity D5000 ScreenTape | Agilent Technologies | 5067-5592 |
| High Sensitivity D5000 sample buffer | Agilent Technologies | 5190-7745 |
| High Sensitivity D5000 ladder | Agilent Technologies | 5190-7747 |
| NovaSeq 6000 S4 Reagent Kit v1.5 (300 cycles) | illumina | 20028312 |
| UltraComp eBeads™ Plus Compensation Beads | ThermoFisher | 01-3333-42 |
| **Deposited data** | | |
| Raw scRNA-seq data | This paper | SRA:PRJNA1090143 |
| Proscessed scRNA-seq data | This paper | GEO:GSE262140 |
| **Experimental animals** | | |
| Mouse: NSG (NOD.Cg-PrkdcscidIl2rgtm1Wjl /SzJ) | The Jaxson Laboratory | Strain #:005557 (RRID:IMSR_JAX:005557) |
| Mouse: NOG-W41(NOD.Cg-Prkdcscid Il2rgtm1Sug Kitem1(V831M)Jic/Jic) | Central Institute for Experimental Animals | N/A |
| Macaca fascicularis | Research Center for Animal Life Science, Shiga University of Medical Science | N/A |
| **Software** | | |
| FlowJo (version 10.8.1) | BD Biosciences | https://www.flowjo.com/solutions/flowjo |
| Python (version 3.8, 3.9, 3.10) | Python Programming Language | https://www.python.org/ |
| R (version 4.3.2) | The R Project for Statistical Computing | https://www.r-project.org/ |
| ELDA (version 1.5.0) | Hu et al. | https://search.r-project.org/CRAN/refmans/statmod/html/elda.html |
| Cell Ranger (version 7.0.0) | 10x Genomics | https://support.10xgenomics.com/single-cell-gene-expression/software/pipelines/latest/what-is-cell-ranger |
| Scanpy (version 1.9.5) | Wolf et al. | https://scanpy.readthedocs.io/en/stable/ |
| DoubletDetection (version 4.2) | Adam et al. | https://github.com/JonathanShor/DoubletDetection |
| scVI (version 1.0.4) | Lopez et al. | https://yoseflab.github.io/software/scvi-tools/ |
| pytorch (version 2.1.3) | Paszke et al. | https://github.com/pytorch/pytorch |
| jax (version 0.4.23) | James et al. | https://github.com/google/jax |
| fa2 (version 0.3.5) | Mathieu et al. | https://github.com/bhargavchippada/forceatlas2 |
| Monocle 3 (version 1.3.1) | Cole et al. | https://cole-trapnell-lab.github.io/monocle3/ |
| ClusterProfiler (version 4.10.0) | Guangchuang et al. | https://guangchuangyu.github.io/software/clusterProfiler/ |
| diffxpy (version 0.7.4) | David et al. | https://github.com/theislab/diffxpy |
